## Supplementary figures and images for "Control of telomere length in yeast by SUMOylated PCNA and the Elg1 PCNA unloader"

### Suppl. Figures

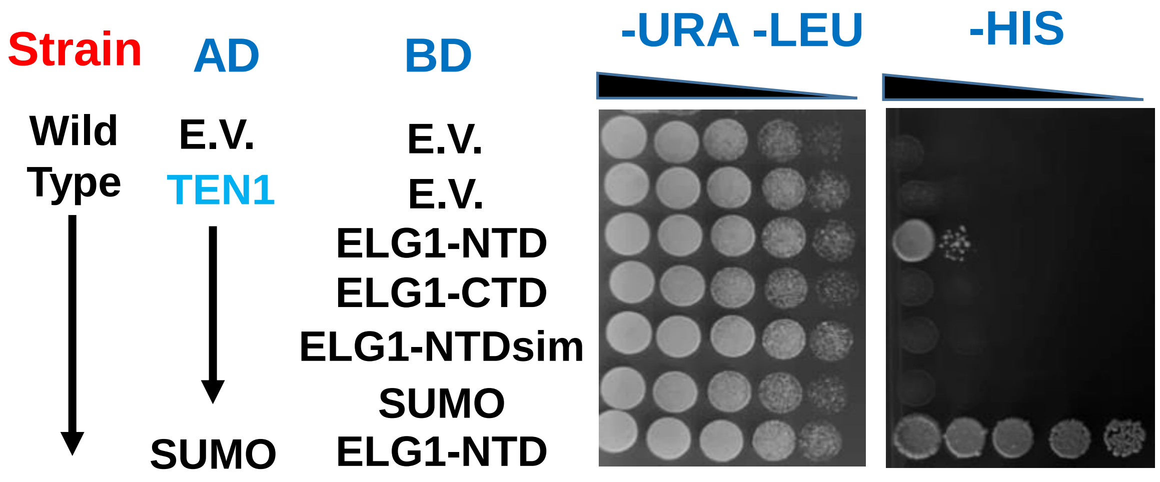

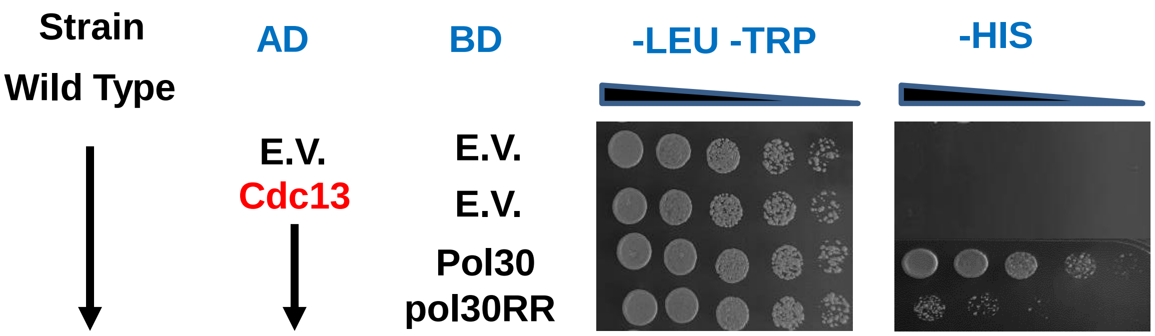

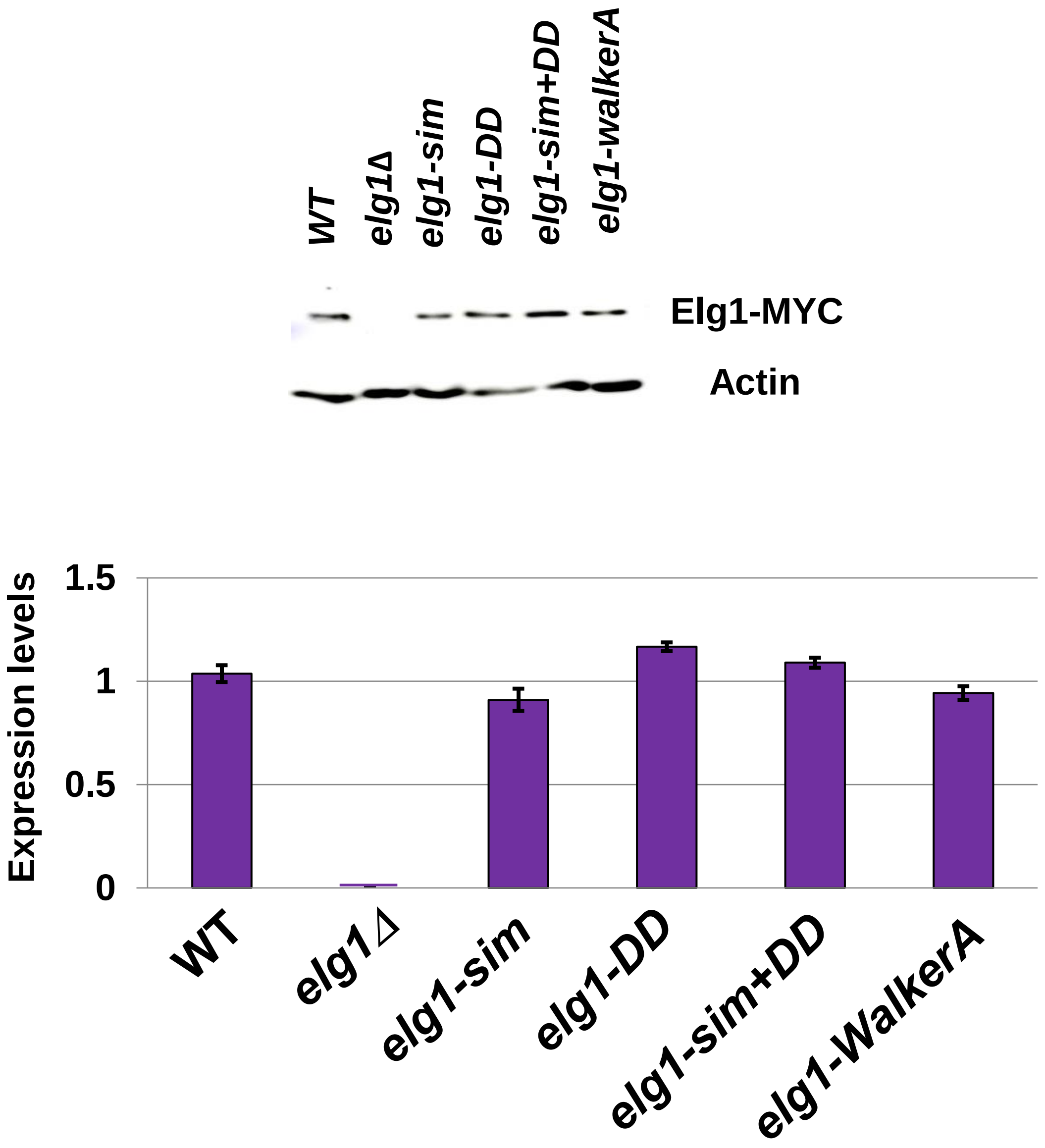

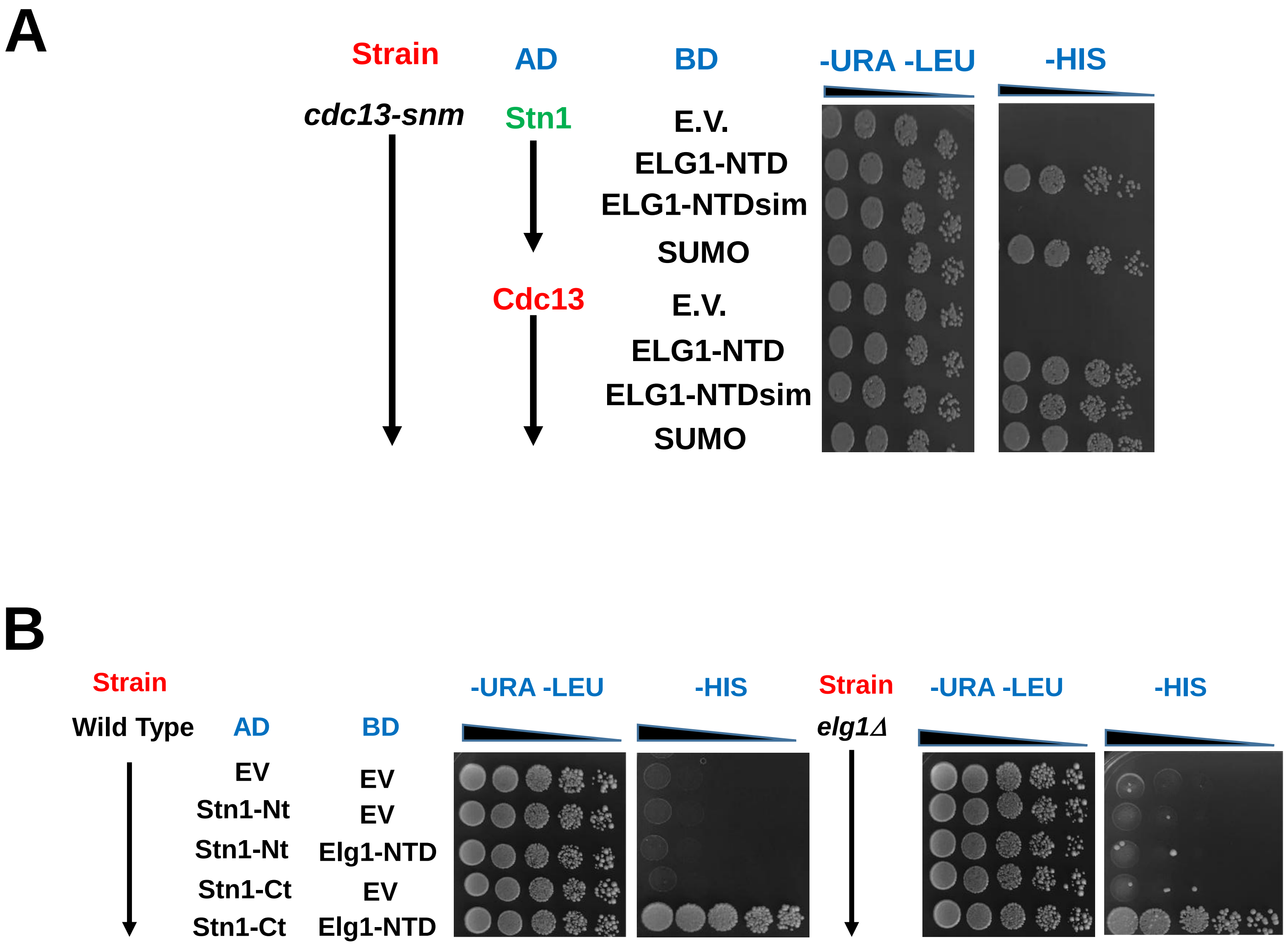

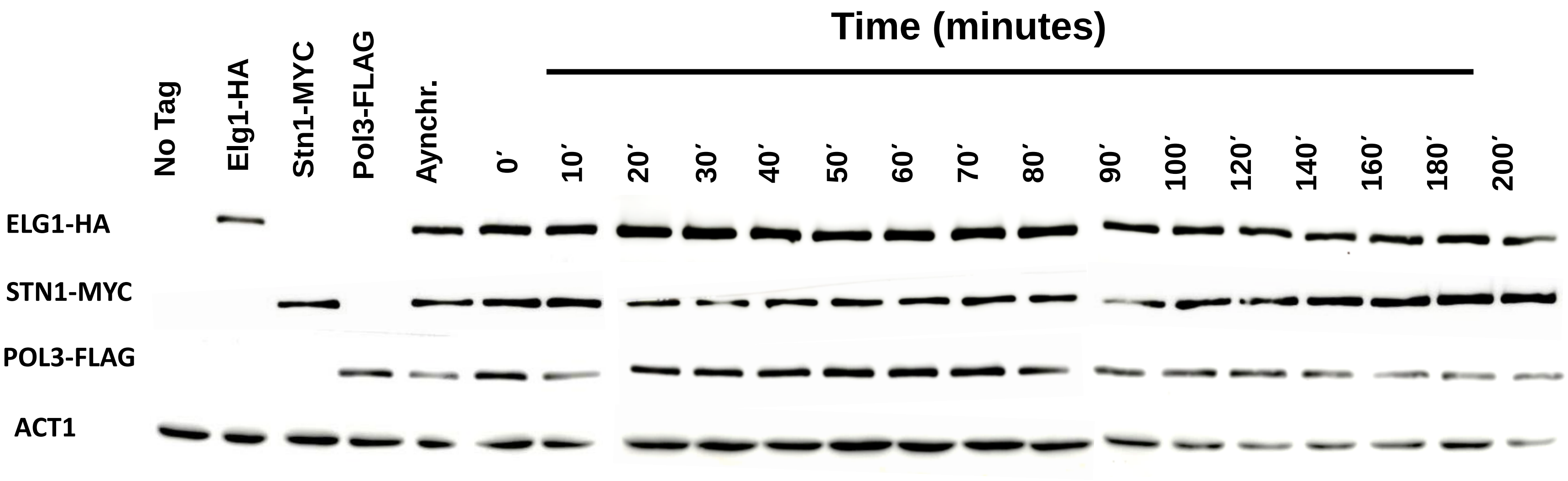

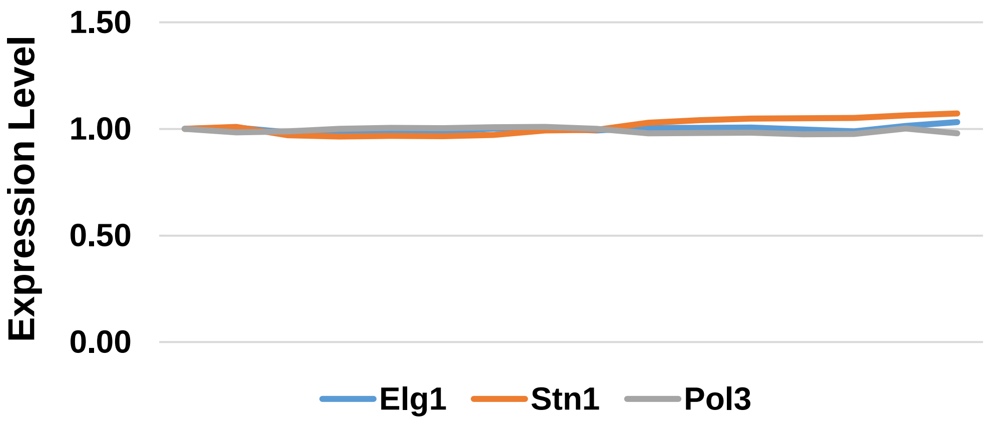
